# Non-invasive brain stimulation protects cognitive impairment in i.c.v.STZ injected rats: role of adult neurogenesis

**DOI:** 10.1101/2025.07.21.666022

**Authors:** Aasheesh Kumar, Avishek Roy, Vedant Karaddi, Suman Jain, Jatinder Katyal, Yogendra Kumar Gupta

**Affiliations:** Department of Physiology, All India Institute for Medical Sciences, New Delhi; Department of Pharmacology, All India Institute for Medical Sciences, New Delhi; division of Clinical Geriatrics, Center for Alzheimer Research, Department of Neurobiology, Care Sciences and Society, Karolinska Institutet, Stockholm, Sweden

**Keywords:** Extremely low frequency magnetic field (ELF-MF), Alzheimer’s disease (AD), adult neurogenesis, i.c.v streptozotocin, dentate gyrus, neuroinflammation, repetitive transcranial magnetic stimulation (rTMS)

## Abstract

In Alzheimer’s like dementia, neurodegeneration and synaptic dysfunction is known to the critical player in explaining the cognitive impairment. Adult neurogenesis, normally a chronic and quite process is explained to have potential in the field of Alzheimer’s therapy. Previous research on the non-invasive brain stimulation showed that controlling pattern of stimulation externally we can regulate/ entrain neuronal activity, possibly altering the structural changes in the circuit level. However, literature investigating if non-invasive brain stimulation could hold any potential to initiate process of adult neurogenesis are scarce. In the present study, with the use of behavioural, microscopic and biochemical tools we found that extremely low frequency magnetic field at an intensity of 17.96µT with sinusoidal wave of 50Hz for 2hr daily for a period of two week in streptozotocin induced animal model of sporadic Alzheimer’s disease can cause improvement in spatial and reference memory, influencing their swimming strategy in water maze. Which is caused by stimulation in immature neural pluripotent stem cells, with additional redox balance and mitigation of glial aggravation in brain areas like olfactory bulb, prefrontal cortex and hippocampus. These changes were accompanied by neuroprotection as observed in the granular layer of dentate gyrus. Taken together, present study explains plausible mechanism of action on protection of neurodegeneration in Alzheimer’s disease through non-invasive brain stimulation.

**Graphical abstract:** Graphical illustration representing external magnetic field causing cognitive improvement eliciting adult neurogenesis in animal model of Alzheimer’s disease

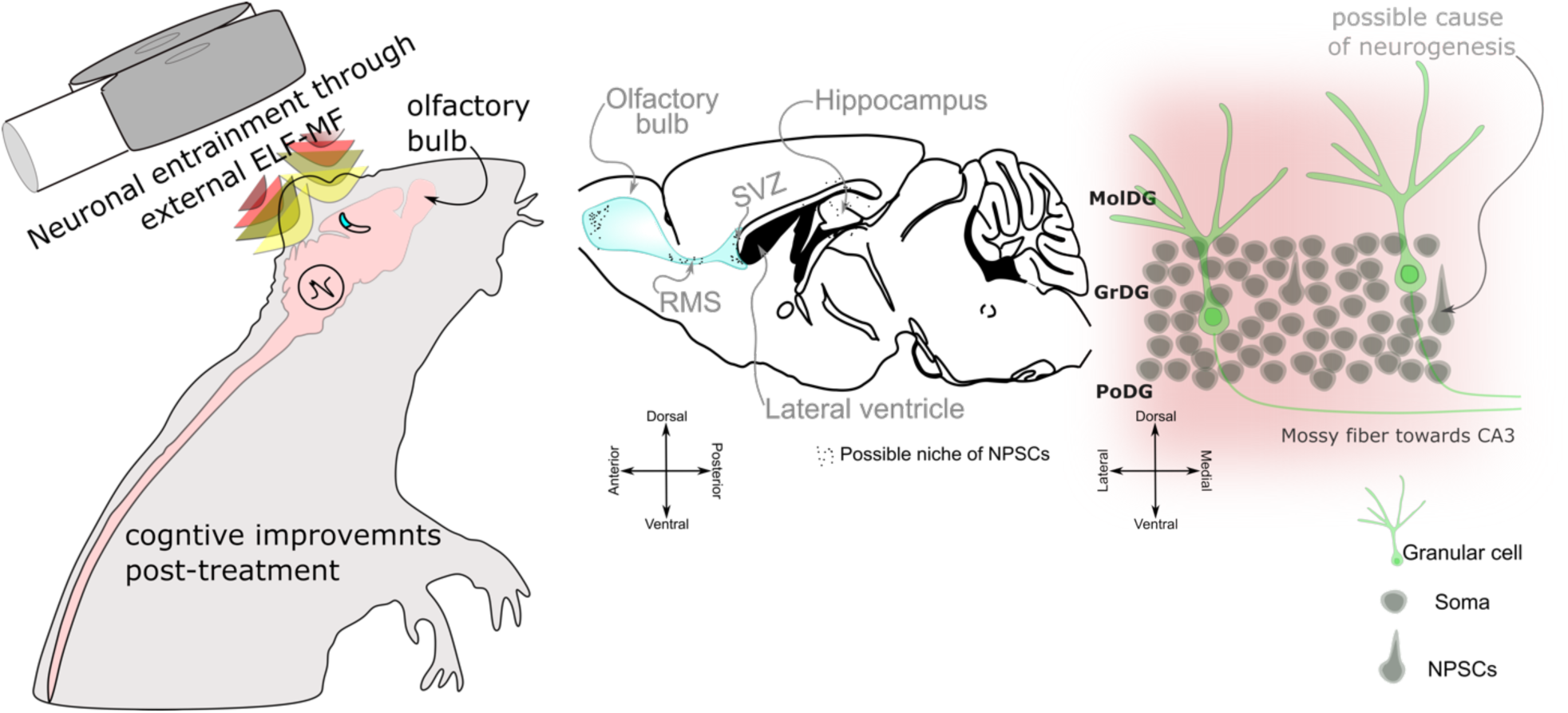

## Introduction

Alzheimer’s disease (AD) is a multifactorial disorder affecting the geriatric populations, which is characterized by deposition of amyloid plaque (Aβ) and neurofibrillary tangles (NFTs) in the central nervous system, with a progressive loss of cognitive abilities (Breijyeh *et al*., 2020; Zhang *et al*., 2024a). One of the early changes that have been reported using the imaging as well as fluid biomarker of AD is loss of insulin signalling and glucose metabolism in brain or in serum (Mosconi *et al*., 2010; Trueba-Sáiz *et al*., 2013). Intracerebroventricular injection of streptozotocin (i.c.v. STZ) can cause Alzheimer’s type dementia with impairments in synaptic dysfunction, cognitive, anxiety, imbalance in oxidative balance, neuroinflammation even imbalance in insulin signaling (Duelli *et al*., 1994; Mehla *et al*., 2013; Salkovic-Petrisic *et al*., 2013; Knezovic *et al*., 2015; Rostami *et al*., 2017; Bassani *et al*., 2018a; Song *et al*., 2018; Roy *et al*., 2022). Previous studies have shown that i.c.v. STZ injection has varied trajectories depending upon the terminal point studied. With a rigorous search on the same line, we found that the first impact on the neuronal as well as behavioural effects were reported after 14 days of incubation period (see supplementary **Fig. 2. b**). In a previous work we have found loss of dendric spine in granular cells of dentate gyrus of hippocampus (Roy *et al*., 2022).

Though the synaptic loss is found to be primary to neuronal degeneration in clinical post-mortem studies in AD (Kumar *et al*., 2024). However, there are less studies exploring what happens to the neuron’s endogenous regenerative property of the neuronal pluripotent stem cells (NPSCs) in AD conditions. Available literature pointed out a sharp drop in the adult hippocampal neurogenesis, a physiological property to generate and incorporate new neurons in the circuitry of AD patients. More specifically these decreases are tightly associated with the cognitive status of the AD patients (Mu & Gage, 2011; Moreno-Jiménez *et al*., 2019; Llorens-Martin, 2020; Salta *et al*., 2023). Interestingly, in 2017 male Wistar rats were studied for adult neurogenesis after 21 days of i.c.v. STZ treatment. Results revealed in spatial, episodic as well as in fear memory with an increase in anxiety, which is accompanied by neuroinflammation and decrease in adult neurogenesis markers like NeuN, DCX, KI-67 as well as proliferation marker BrdU (Bassani *et al*., 2018b).

To settle down to the marker for neurogenesis to be studied here, we rigorously checked the mammalian adult neurogenesis gene ontology database (Overall *et al*., 2012a). After the outcome/ process and cell stage filtration we found one of the markers i.e. *nestin* for immature neurons, which indicate either ‘determined progenitor’ or ‘stem cell’ as the stage of NPSCs development (see **Fig.** 3.e.i-iii.).

In the AD therapeutics available with anti-cholinergic/ NMDAr blocker drugs, or clinical trials are going on immune therapy (Jicha *et al*., 2021; Zhang *et al*., 2024b). However, most of the studies land up to failure. Currently there are numerous non-pharmacological modalities available for AD pathology from invasive protocols like deep brain stimulation to non-invasive stimulation like photo-biomodulation, ultrasound and transcranial magnetic field stimulation (Ning *et al*., 2022). Though the working principles are different, the idea of non-invasive brain stimulation was to entrain neuronal activity to the external pattern of stimulation. In humans, repetitive transcranial magnetic field stimulations (rTMS) at dorsolateral prefrontal cortex shown to have cognitive improvement in AD patients with anti-AD drug or even without any drug (Hallett, 2000; Cotelli *et al*., 2008, 2011; Wei *et al*., 2023). Similar outcomes were also reported in transgenic mice model of AD with alteration in cortical long-term potentiation (LTP), excitability as well as spatial learning and memory (Wang *et al*., 2015). In another study rTMS treatment in animals at <1Hz i.e. low frequency led to CaMKII mediated synaptic plasticity in hippocampus (Li *et al*., 2019), similarly in BL6 mice administered cuprizone (per orally) showed a rescue of cortical pathology also production of cytokines after theta burst stimulation (Yang *et al*., 2020). In Aβ_1-42_ induced cognitive deficit animals, low frequency magnetic field stimulation found to have cognitive improvement through replenishing hippocampal neurotrophins and LTP (Tan *et al*., 2013). In another study, ELF-MF-stimulation in Gerbils at 50Hz frequency not only improved neuronal health, but also causes a reduction in the activation of reactive astrocyte as well as microglia in hippocampus (Rauš *et al*., 2012). Further, a study by Choung J.S. et al., 2021 attempted to investigate efficacy of rTMS treatment at both high and low frequency rTMS (20Hz and 1Hz), where only high frequency magnetic field stimulation showed to have enhanced expression of neurogenesis (Choung *et al*., 2021). It is important to note that in this study, authors used 1600 pulses per session with 1.26 T and NeuN and nestin expression is only upregulated in cortical samples not hippocampus. In surgical model of stroke, ELF-MF stimulation at 1mT with 50Hz frequency for 28 days showed gain of function in BrdU/NeuN^+^ cells in sub-granular cells of hippocampus that are accompanied by Notch 1, Hes 1, and Hes 5 increased expression (Gao *et al*., 2021a).

Taking together, in the present study we set out to investigate the influence of the early i.e. stem-cell (nascent) stage NPSCs of dentate gyrus, after one-week low frequency magnetic field stimulation at 17.96µT in modified Helmholtz coil in the Wistar rats intracerebroventricularly injected with streptozotocin (3mg/kg BW; bilateral injection). We purposefully started BrdU injections in rats only after a week of ELF-MF stimulation, in order to mark especially day of non-invasive stimulation. Further, we studied spatial and reference memory complemented with oxidative stressor, neuronal health and neurogenesis using standard behavioral, biochemical and histological tools.

## Methods and materials

### Subjects and animal group division

39 adult female albino Wistar rats weighing 220±10 gm was used in the study. Animals were provided standard laboratory food pellets (Ashirwad Co. India) and RO water *ad libitum*. Furthermore, room temperature of 25± 2°C and light: dark cycle of 14:10h and 50-55% relative humidity were maintained. All the animals were housed in separate polypropylene cages (50 cm X 20 cm X15 cm) with metallic lid, which had separate laboratory chow and water bottle holders. All the experimental procedures are approved by institutional animal ethical committee vide no. 12/IAEC/2017. This was in accordance with the committee for the purpose of control and supervision of experiments on animals (CPSCEA) guidelines.

Rats were randomly divided into three groups, namely i.c.v.aCSF, i.c.v.STZ and i.c.v.STZ+MF. Each group of animals consists of 9 animals. 0.12% of animals were excluded due to death before completion of the study period. i.c.v.aCSF treated groups of animals received 5µl bilateral artificial cerebrospinal fluid (aCSF) in the lateral ventricles bilaterally and then exposed to sham ELF-MF i.e. no stimulation was given to the animals. These animals were kept in polypropylene cages for 2hrs, this counterpart represents the vehicle injected placebo group. In the i.c.v.STZ group animals received 3mg/kg streptozotocin dissolved in aCSF in the lateral ventricles with 5µl/site bilaterally. On the third group i.e. i.c.v.STZ+EMF i.c.v. was STZ administration was common, but these animals were further exposed to the ELF-MF (17.96µT, 50Hz, 2hr/day) in the magnetic field chamber (see supplementary fig.1). For behavioral studies 9 animals/ group were used, whereas for oxidative stressors and histological examinations 4, 5 animals/ group were used respectively.

**Figure 1.:**
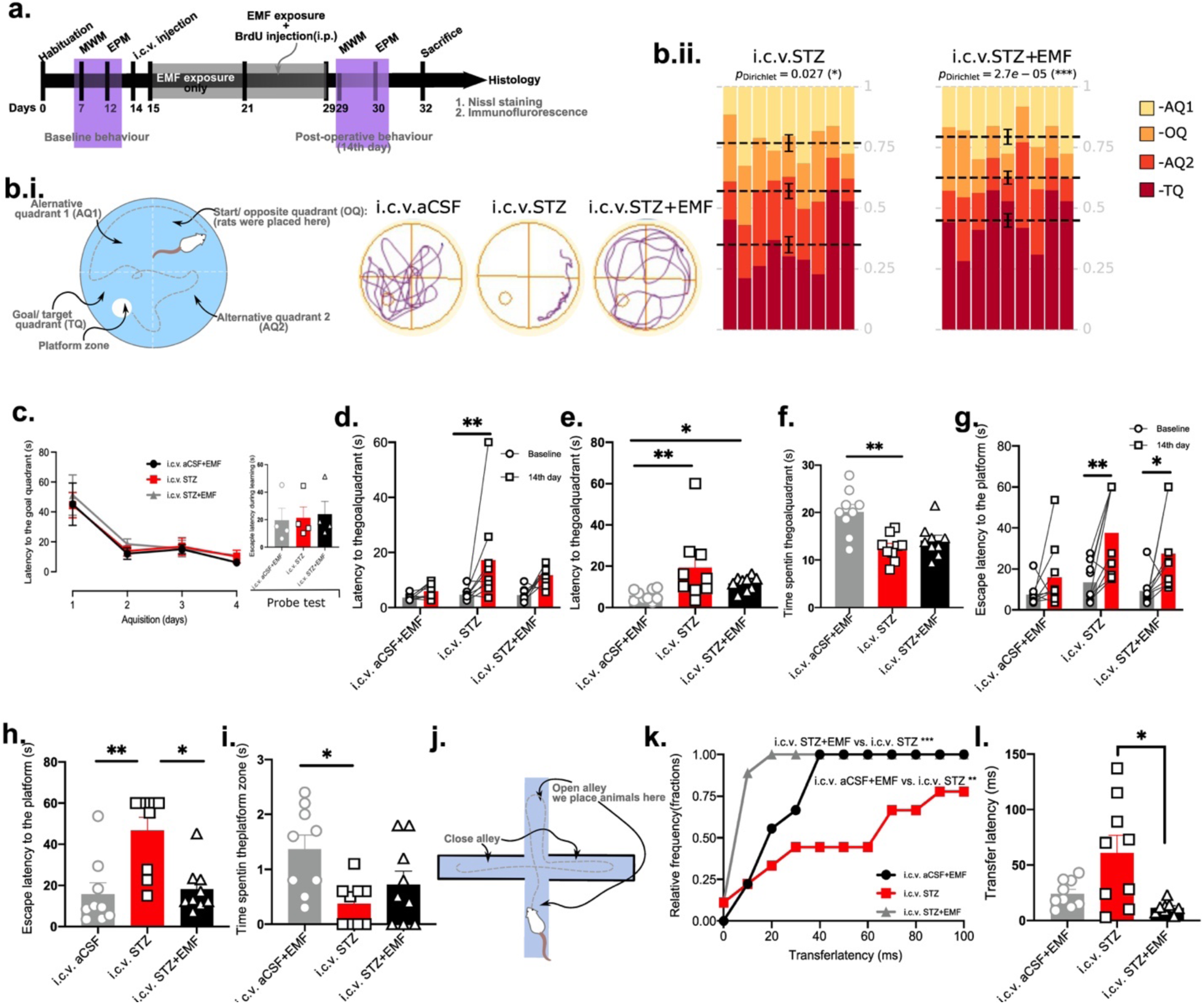
Cognitive improvement after non-invasive brain stimulation; illustration showing study design with chronological order which encompasses 32 days in total from the day of habituation, purple shades represents the behavioural investigations, whereas dark shade represent the ELF-MF exposure period (a); illustration of MWM setup with the imaginary zones used in the experiments with representative track-plot of the swimming (note the quadrant codes, b.i.); distribution of fraction of time spent in four quadrants using likely-hood ratio statistics using Dirichlet distribution in disease and treatment group (b.ii.); escape latency of the animals investigated on the terminal phase for understanding learning, where inset histogram represent the average escape latency only on the probe trial of the post-operative MWM test (c); parameters from goal quadrant latency between pre- and post-operative days within the group (d), latency to the first entry and time spent in goal quadrant on post-operative day probe trial (e-f); parameters from platform zone latency to the first entry between pre- and post-operative days within the same group (g), latency and time spent in platform zone on the post-operative day between three groups (h-i); illustration of the elevated plus maze protocol being used in the present study to understand reference memory (j); relative frequency of transfer latency and average transfer latency as measured in elevated plus maze after 14 day post-operative to STZ injection (k-l); data represented as mean±SEM; for escape latency for four days 14^th^ day post-operative to i.c.v. STZ injection and relative frequency of transfer latency plotted as line-graph whereas other data has been plotted as histogram with individual data points; p≤0.5 is considered as significant; one-way ANOVA for three separate group comparison, two-way ANOVA for escape latency for four days at the terminal behavioural examination, Kolmovogorov-Smirnov test for relative frequency was adapted whereas for normality check we’ve used Shapiro-Wilk test; AQ1= alternative quadrant 1, AQ 2= alternative quadrant 2, OQ= opposite quadrant, TQ= target quadrant.

### Study design

After adaptation to the departmental animal facility, rats were randomly divided into three groups using block-size randomization protocol (Efird, 2011). Animals were quarantined in the departmental animal facility, when they were checked for any infections, if any animals were provided with gentamycin for 3 days. And we have used this period to acclimatize to the polypropylene boxes used as restrainer during ELF-MF exposure.

All the baseline behavioral recordings were performed after the quarantine period of 7 days. After 5days of MWM test on 12^th^ day EPM was used for reference memory assessment. After 2 days animals were either injected with aCSF or STZ intracerebroventricularly through stereotaxic manipulations. After one day of recovery period from surgery, only i.c.v.STZ+EMF group were exposed to magnetic field stimulation for 14 days. During this period simultaneously i.c.v.aCSF group were kept in the polypropylene boxes for 2hr duration/ day for 14 days. Which act as sham-stimulation proxy in the study. All the animals were intraperitoneally injected with 5-bromo-2’-deoxyuridine (BrdU) at a dose of 50mg/Kg for 7 consecutive days along with continuation of ELF-MF exposure. On 29^th^ day animals were finally tested for change in spatial memory and its learning and reference memory to assess terminal behavioural changes post-operative 14^th^ day. Finally, animals were sacrificed and tissues for biochemical (i.e. fresh tissue snap frozen) and histological (i.e. fixed with transcardial perfusion) experiments are collected (**Fig.** 1.a).

#### Behavioral tests

In the present study, Morris water maze (MWM), elevated plus maze (EPM) were chosen to assess alterations in spatial memory, reference memory respectively. All behavioral assessments were recorded and analyzed with video tracking devices. Behavioral changes were recorded on baseline i.e. prior to any i.c.v. injection and then followed on 14^th^ day post-injection of aCSF/STZ alongside ELF-MF exposure on respective groups.

### Morris water maze test for evaluation of spatial memory

We have followed procedures for the MWM test previously described (Morris, 1984; Roy *et al*., 2022). The MWM tank was customized to 170cm in diameter with 72cm in height made from stainless steel. This tank was divided into four equal imaginary quadrants in the video tracking software (AnyMaze ver.05; Stoelting Corp., USA). Animals were trained to remember how to escape swimming by entering a hidden platform. Hidden platform size was 35cm in height and 10cm in diameter for holding animals. Always trials were started by placing animals facing a tank wall, trial was terminated once animal sit on the platform for 20 seconds. If 120 seconds elapsed without finding the hidden platform, then the animal were hand guided by the experimenter. Four days, four trials/day for 2 minutes were spent for training period, on fifth day a probe trial was conducted without hidden platform in the tank. On probe trial day animals got only 60 seconds to perform/ search for the platform zone coordinate (**Fig.**1b.).

On the 14^th^ day post-injection, four-day training protocol was performed to understand the effect of ELF-MF on learning curve of spatial information.

### Elevated plus maze test for evaluation of reference memory

The elevated plus maze (EPM) tool was used to assess reference memory like behavior explained previously (Itoh *et al*., 1990; Sharma & Kulkarni, 1992). This behavior is based on the approach-avoidance conflict in rodents in response to elevation. Animals have a natural tendency to escape from an open alley of EPM to enclosed alley, it is noted that after one exposure to EPM, the latency to escape from open alley to enclosed alley is reduced. However, this phenomenon is dependent on the time spent on enclosed arm on 1^st^ day. For the present study we only used animals with closed arm duration >40s.

The length of the open and closed arm of 76 and 134 cm, respectively, with a junction of arms, 30 i.e., central area of 6 × 6 cm with an elevation height of 45cm from the ground. Behavior was video tracked with CCD camera (Panasonic, Japan) associated with Ethovision XT (Ver. 07, Noldus, Netherlands). Rats were placed individually at the end of either of the open arms and the duration animal took to move from open to enclosed arm (TL) was noted on the 1^st^ day. The animals were allowed to explore the apparatus for 60s. On the next day, after the first exposure TL was again noted. Second day TL is plotted as a measure of reference memory on baseline as well as 14^th^ day post-injection (**Fig.**1a;j). The apparatus was properly cleaned between two trials to avoid influence of previous animals.

#### Intracerebroventricular injection

In the present study, we have injected artificial cerebrospinal fluid (aCSF) prepared based on recipe from (Mehla *et al*., 2012) with an osmolarity of 295±5 mmol/kg or streptozotocin (STZ, 572201-1GM, Merck Milipore, USA) at a dose of 3mg/kg of BW bilaterally with a volume of 5µl/ site (same for aCSF). Stereotaxic surgical procedure, selection of dose and coordinates are explained previously (Roy *et al*., 2022). The coordinates for the stereotaxic injection were AP: - 0.84 mm; ML: ±1.5 mm; DV: −3.5 mm under thiopentone anesthesia (50mg/Kg body weight), with glycopyrolate (i.p., 80µg/kg of body weight) as anti-cholinergic drug to avoid choking during surgery following the rat atlas (Paxinos & Charles Watson, 2007).

#### Magnetic field stimulation

In order to study the effects of extremely low frequency magnetic fields (ELF-MF) we have exposed the animals to a modified Helmholtz coil, which consists of four circular coils. Two outer coils consist of a coil turn ratio of 18, whereas two inner coil turn ratio was 8. Diameters of coils are1,000 mm, with 45 mm width. Distance of outer coil from center of the structure is 470 mm and of the inner coil is 122 mm. Current in electromagnetic coil is 1 A at 50 Hz. Magnetic field is 17.96μT at central exposing/ platform area (see supplementary **Fig.**S1 for more details).

Animals were exposed to the magnetic field for two weeks post-injection of STZ where they were given 2hr/ day ELF-MF stimulation at a frequency of 50Hz, which was regulated through stabilized current supplier (**Fig.**1a).

### Marking proliferative cells with 5-Bromo-2’-Deoxyuridine (BrdU)

We have used BrdU (19-160, Sigma Aldrich), a thymidine analogue that can be incorporated at S-phase of cell proliferation, to look at granular cells at dentate gyrus with proliferation/ in DNA duplication phase dissolved in normal saline (0.9% NaCl) as mentioned elsewhere (Wojtowicz & Kee, 2006). As stated, that our goal was to tag cells specifically after one week of ELF-MF exposure, hence we have started BrdU injection (i.p., once/day, 50mg/kg body weight) only after one week of ELF-MF exposure. Which was carried to 28^th^ day (**Fig.**1.a).

BrdU was injected immediately after ELF-MF exposure for 2hr. And then animals were monitored for any abnormal behaviors in the home cage.

#### Animal sacrifice

At the end of the study, either the animals were sacrificed through CO_2_ concussion for analysis of oxidative stressors in hippocampus and frontal cortex isolated separately, or transcranial perfused after lethal dose of sodium thiopentone (150mg/Kg of BW) accordingly.

### Tissue preparation for biochemical analysis

After CO_2_ concussion, the brain was isolated followed by decapitation and immediately it was placed on to the ice-cold PBS (0.1M) to isolate hippocampus and frontal cortex carefully with no contamination from other nearby tissues. Next, brain areas from both contralateral hemispheres were pooled together for each animal. 5 animals/ group were used for the measurement of SOD1 activity and GSH. After taking the net weight of the samples, it was diluted 10 times with 0.1 M PBS. Then, tissues were immediately homogenized using Homogenizer (Gentle MACS Dissociator, Miltenyi Biotec). Samples were then centrifuged at 5000 rpm for 10 minutes at 4°C. Supernatant was collected and used for the biochemical assay. Reading of 96-well plate and cuvette for reaction mixture of GSH and SOD assay was taken using *Gen5 3.03 software* of Spectrophotometer (Epoch2 Microplate reader, BioTek).

### Measurement of Superoxide dismutase 1 (SOD1) activity

The estimation of SOD1 activity was carried out by the method of (MARKLUND & MARKLUND, 1974) with slight modification, utilizing the inhibition of auto-oxidation of Pyrogallol by SOD1 enzyme. Final reaction mixture was made 3 mL with distilled water. Reaction mixture contains 1.5 mL of Tris Buffer (100 mM, pH = 8.2), 0.15 mL of Pyrogallol (0.2 mM in 0.5 N HCl) and 10 µL of brain homogenate. Control reaction mixture was devoid of brain homogenate. Auto-oxidation of Pyrogallol under alkaline condition generates free radicals which are used by SOD1 enzyme present in brain lysate. Decrease in auto-oxidation shows indirect evidence of SOD1 activity. Spectrophotometric reading was taken at 420 nm for 180 s after each 10 s interval.

### Measurement of reduced glutathione (GSH) level

Level of Glutathione was estimated by using the method of (Moron *et al*., 1979). First, glutathione standard (0.1mg/mL) was used to plot a standard curve. Then the concentration of glutathione in the sample was estimated using a standard curve. The principle behind the glutathione assay utilizes the color reaction for 0.04 % 5,5’-dithiobis-(2-nitrobenzoic acid) or DTNB in the 0.1 M Phosphate buffer (pH=8) and 10 % Trichloroacetate (TCA). The optical density was taken at 412 nm wavelength after the incubation of reaction mixture for a maximum of 10 minutes.

### Transcardial perfusion and tissue harvesting

Under lethal dose of sodium thiopentone (150mg/kg body weight, i.p.), without any anti-cholinergic drug. Animals were first confirmed for deep anaesthesia with paw-pressure test. animals were placed onto an icepack on its back and immediately a lateral incision was made in the skin and abdominal wall and then small incisions were made to access the plural cavity by through incising diaphragm and cardiac envelop was excised to access the heart (Gage *et al*., 2012). Finally, preceding 0.9% ice-cold NaCl solution paraformaldehyde (4%, in PBS, W/V) was perfused <50ml each animal. Next, brain samples were carefully isolated without physical damage and post-fixed in the same solution for 2-3 days before further processing.

Next, 20µm thick coronal 6^th^ alternative sections were cryo-sectioned in cryotome (HM550, Microme, Thermo Fisher) after cryoprotection with graded (15%, 30%) sucrose solution in PB (W/V) and taken onto gelatin (3%, W/V) coated cloned frosted glass slides (75X25mm; BlueStar Pvt. Ltd.). Sections were kept in −80°C until further use.

### Histology

The brain sections were stained with cresyl violet (0.4%) for 3 min, dehydrated and mounted with DPX. For comparison of hippocampal sections, we restricted to AP-axis from −3.80 to −3.92 mm (**Fig.** 2.e)in dentate gyrus (DG) layers like molecular layer of dentate gyrus (molDG), granular layer of dentate gyrus (grDG) and polymorph layer of dentate gyrus (poDG) with guidance from atlas (Paxinos & Charles Watson, 2007). The stained sections were kept in the dark until imaging.

**Figure 2.:**
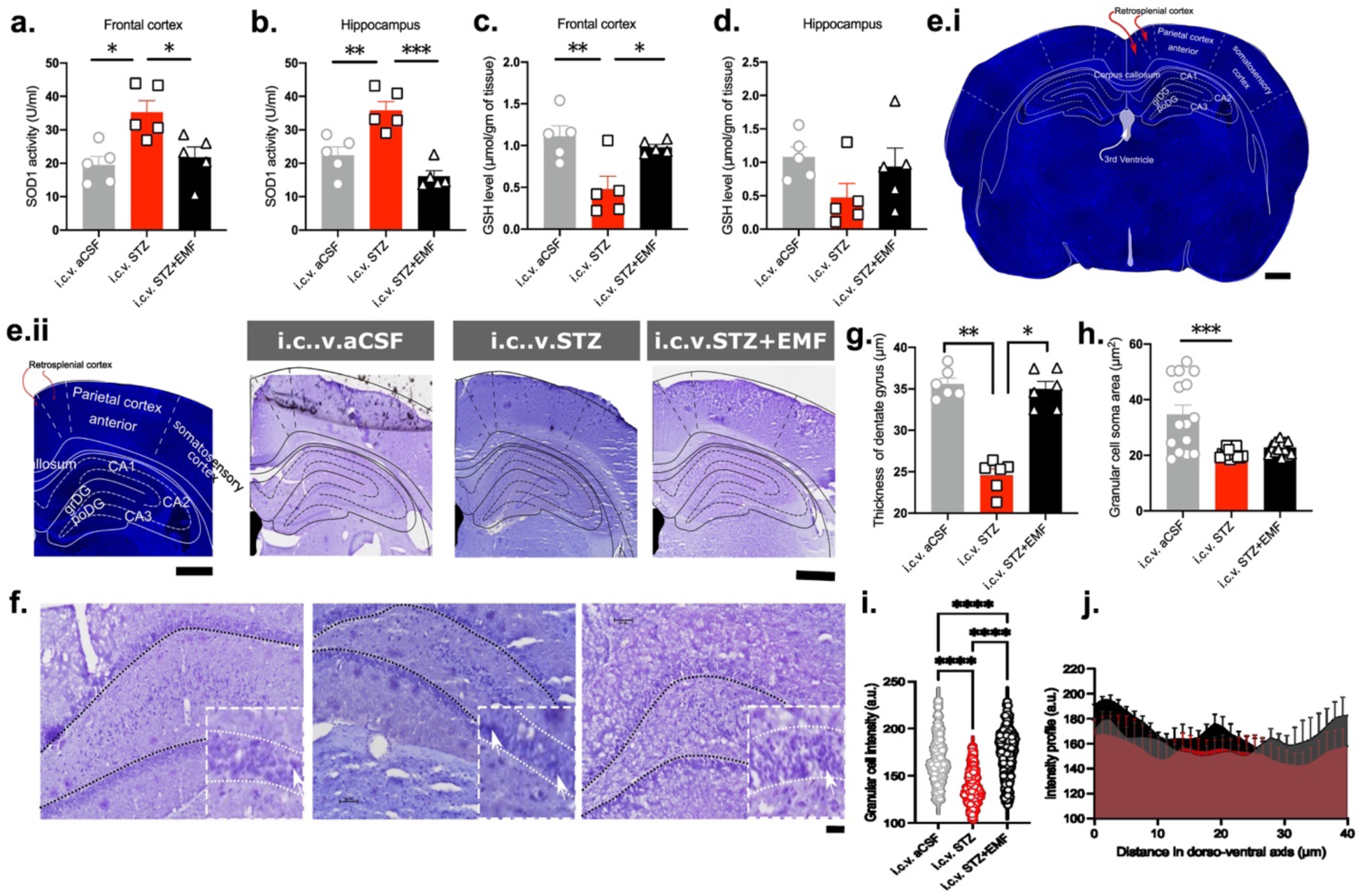
Non-invasive brain stimulation causes improvements in redox balance and repairing the structural loss induced by STZ; alteration in the superoxide dismutase activity and reduced glutathione level in the frontal cortex and the hippocampal homogenates after 14 day ELF-MF exposure in the STZ injected rats (a-d); illustration to represent the coronal section of rat brain stained with DAPI, how the anatomical landmarks are being used for immune and cytoarchitectural examination in the present study (ei); panel of the represented coronal sections stained with DAPI with landmarks and CV stained sections from i.c.v.aCSF, i.c.v.STZ and i.c.v.STZ+EMF group (from left to right, eii); represented images of dentate gyrus with zoomed images in the inset (arrow marked pyknotic cell with pale soma, f); thickness of the granular layer of DG and soma area of granular cells of DG (g-h); intensity of granular cells as a proxy of pale-ness due to chromatolysis (j): intensity profile of DG layer across dorso-ventral axis (k); data is represented as histogram with individual data points; n=5 for biochemical examinations and n=3 for histological examinations with different fields at high magnification; p≤0.5 is considered as significant; one-way ANOVA was used for the statistical comparison between three groups whereas for normality check we’ve used Shapiro-Wilk test; scale = 50 micron for panel ‘e’ and 100 micron for panel ‘f’ images.

#### Immunostaining

Sections were permeabilized with PBST (0.2% Tween-20 in PBS), blocked in 5% BSA for 2 hr at room temperature before being incubated in anti-BrdU (1:200; MAB3424, Sigma Aldrich, raised in mouse), anti-Nestin (1:100; N5413, Sigma Aldrich, raised in rabbit), anti-GFAP (1:500; G3893, Sigma Aldrich, raised in mouse), anti-IBA-1 (1:50; Kab04096, Kinesis, raised in rabbit) for 2 days at 4°C. Next, sections were incubated with secondary antibodies (anti-rabbit, 1:400; DI-1594; anti-mouse, 1:1000; fl-2000; Vector laboratories, USA) for 2 hr at room temperature. After washing, sections were mounted with fluoroshield with DAPI (F6057, Sigma-Aldrich, USA) and imaged using fluorescence microscope.

For BrdU and Nestin co-localization we have used protocol previously described (Wojtowicz & Kee, 2006).

#### Image acquisition and image analysis

Brain sections stained with cresyl violet were imaged using the brightfield Eclipse Ni upright microscope. For fluorescence imaging of the sections probed with primary antibodies, same microscope was used additionally with fluorescence lamp 49 (C-HGFI, Nikon, Japan) using triple filter block of DAPI (blue; excitation at 385-400 nm; bandpass at 393CWL), FITC (green; excitation at 475-490 nm; bandpass at 483CWL) and TRITC (red; excitation at 545-565 nm; 51 bandpass at 555CWL), according to the need. Images were captured using a color camera (DSRi-2, Nikon; 52 Japan) and saved in RGB format. We have used 4 animals/ group and 4 sections/ animals with high magnifications either 20X or 40X objectives.

Images of cresyl violet we analyzed for layer thickness and granular cell soma volume following usual histometric procedure using Fiji (Schindelin *et al*., 2012). For BrdU/Nestin dual positive colocalized sections have been used ‘multipoint tool’ for counting the number in each field of view. However, for more detailed analysis of the sections from i.c.v.STZ+MF group we have used ‘just another co-localization plugin’ and plotted the cytofluorometry results between BrdU vs. Nestin signals for Pearson correlation (Bolte & Cordelières, 2006).

### Cytofluorometry for colocalization analysis of BrdU and nestin signals

For colocalization measurements, we have calculated both the dual positive cells in same section probed with anti-BrdU and anti-nestin separately as well as combined using the ‘threshold color’ option in Fiji (Schindelin *et al*., 2012). Then, interaction between the signals from BrdU vs nestin fluorescence intensity was further looked under Pearson’s correlation with Manders’ overlap coefficient, which gives the influence of one signal on the other channel co-localized with it using JACoP plugin (Bolte & Cordelières, 2006). Cytofluorogram data were further processed for joint plot using seaborn library.

### Swimming strategy distribution of animals during spatial memory task

In order to understand if there is any alteration in the swimming strategy in the animals underwent STZ treatment and STZ treatment along with magnetic field stimulation, we performed likely-hood ratio of uniformity distribution of the time-spent in the four quadrants. To do so we’ve employed Dirichlet distribution statistics explained elsewhere (Maugard *et al*., 2019). Four quadrants were conceptualized as one target quadrant (TQ), one opposite quadrant (OQ), and two alternative quadrants (AQ). Null hypothesis was adapted only when a group’s fraction of 43 time spent in TQ=AQ=OQ. In the present study we’ve used percentage of time spent data from all the aforementioned quadrants from the probe trial day 14^th^ day post-injection of streptozotocin.

#### Statistical analysis

To understand distribution of data, Shapiro-Wilk normality test was performed. Between group comparison of behaviour on 14th day post-injection, Kruskal-Wallis test with Dunn’s post-hoc test was performed. However, comparison between baseline vs. 14th day post-injection was done using two-way ANOVA (non-parametric, two tailed, ordinary) was performed with Bonferroni’s correction for multiple comparison. Biochemical and histology analysis was performed with ordinary ANOVA with Bonferroni as multiple comparison. Finally, the association of BrdU and nestin signal in the ELF-MF treated group was measured using Pearson’s correlation. All statistical analysis and graphs were plotted in GraphPad Prism 8.0 (GraphPad Software Inc, San Diego, CA) except for cytoflurometry data which was plotted using scatterplot in seaborn library (Waskom, 2021). A p-value of < 0.05 was considered significant and data were shown as individual values with mean and standard error of mean (±SEM).

## Results

### Non-invasive brain stimulation improve spatial and reference memory

After 14 day of extremely low frequency magnetic field stimulation (ELF-MF, 17.96µT, 50Hz, 2hr/ day) escape latency to the goal quadrant did not alter either within the same groups (p=0.987) or between the groups (p>0.999). However, in i.c.v.STZ group there was an increase in latency to the goal quadrant on 14th day post-injection (p=0.006) as compared to its baseline. Further, i.c.v.STZ injection also causes an incease in latency to the goal quadrant to be increase in both i.c.v.STZ group (p=0.008) and i.c.v.STZ+MF group (p=0.031) as compared to i.c.v.aCSF group. Suggesting, an improvement in cognitive performance after 14 days of ELF-MF stimulation, however the changes are higher than sham exposed group. In continuation, we also have checked latency to the platform zone between baseline and 14th day post-injection revealed an increase in both i.c.v.STZ (p=0.005) and i.c.v.STZ+MF (p=0.043) groups. Interestingly, on 14^th^ day the latency to the platform zone in i.c.v.STZ group was found to be increased in comparison to i.c.v.aCSF (p=0.002) as well as i.c.v.STZ+MF (p=0.037) counterpart. Time spent in the platform zone was only found to be reduced in i.c.vSTZ group (p=0.029) in comparison to i.c.v.aCSF counterpart (Fig. 1.c-i).

These changes in MWM was further investigated for any influence in swimming pattern. To do so, we’ve performed likely-hood ratio statistics (LRS) with Dirichlet distribution. On the 14^th^ day post-injection of STZ, Dirichlet distribution of fraction of time spent in each quadrants. In this statistics we considered equal time spent in each quadrant as null hypothesis (H0) and the alternative hypothesis (H1) was that at least one quadrant will differ significantly.Statistics revealed significant uniform distribution in i.c.v.STZ (p= 0.0274877, LRS= 9.13988), however it was more prominent after of magnetic field stimulation to the STZ treated group (p= 2.68702e^-05^, LRS=23.8482). This finding suggests, improvement in the spatial memory in i.c.v.STZ+MF group compared to i.c.v.STZ could be influenced due to alterations in their swimming strategy. Which was even supported by their individual trackplot (**Fig.** 1.bi-ii).

Apart from MWM we have also employed specific reference memory protocol using EPM setup, where relative frequency of transfer latency was found to be increased in both i.c.v.aCSF (p=0.006) and i.c.v.STZ+MF (p=0.0002) as compared to that of i.c.v.STZ. Further, total transfer latency 14^th^ day post-injection of STZ was also fund to be increased as compared to ELF-MF treated group (p=0.015, **Fig.** 1.k-l).

### Acute electromagnetic field stimulation ameliorates oxidative stress in hippocampus and frontal lobe

Superoxide dismutase 1 (SOD 1) activity was found to be increased in STZ injected rats as compared to that of i.c.v.aCSF (p=0.010) as well as i.c.v.STZ+MF (p=0.027) counterparts in frontal cortex. In hippocampus, simillar changes were found where SOD1 activity in i.c.vaCSF+EMF (p=0.004) as well as i.c.v.STZ+MF (p=0.0002) was reduced as compared to STZ injected counterparts after 14 day post-injection (**Fig.** 2.a-b).

Further, GSH level was only found to be significantly reduced in i.c.v.STZ treated rats as compared to i.c.v.aCSF (p=0.007) and i.c.v.STZ+MF (p=0.031) only in frontal cortex. However, in hippocampus no changes (p=0.1649) were observed (**Fig.** 2.c-d).

### Electromagnetic field stimulation causes alteration in the cytoarchitecture of dentate gyrus granular layer in hippocampus

As mentioned in the method we have focused on the alterations in the dorsal hippocampus from –3.80mm to –3.92mm in AP axis. In the fields examined we have majorly focused on the three layers molDG, grDG and poDG (**Fig.** 2.ei). There was gross atrophy of hippocampus as well as cortical tissue in i.c.v.STZ group (supplementary **Fig.** 2.c).

We have found significant thinning of DG layer in the i.c.v.STZ group (p<0.0001) when compared to that of other counterparts. Suggesting a loss of cellular compactness of DG granular cell layer. Further, to understand more histometric alteration specific to the granular cells we have examined the soma volume. We found a significant reduction in soma volume in the i.c.v.STZ group (p=0.0002) compared to i.c.v.aCSF group (**Fig**. 2.g-h).

### Electromagnetic field stimulation improves glial aggravation and elicits adult neurogenesis

Glial fibrillary acidic protein (GFAP) and ionized calcium binding protein (IBA-1) were used to track astrocytes and microglial activation in the dentate gyrus as well as olfactory bulb (supplementary **Fig.** 2.a). We did not only find increased expression of GFAP and IBA-1, but also the primary projections of the astrocytes were found to be thick in i.c.v.STZ group as compared to that of the i.c.v.aCSF and i.c.v.STZ+MF group. We’ve plotted the expression profile across AP axis of brain in cartesian frame. We found gain of expression of both markers were found to be less after two weeks of magnetic field stimulation in all the brain regions examined (viz. Olfactory bulb, hippocampus and frontal cortex) (**Fig.** 3.f.). These results further underpin the potential of extremely low EMF stimulation.

**Figure 3.:**
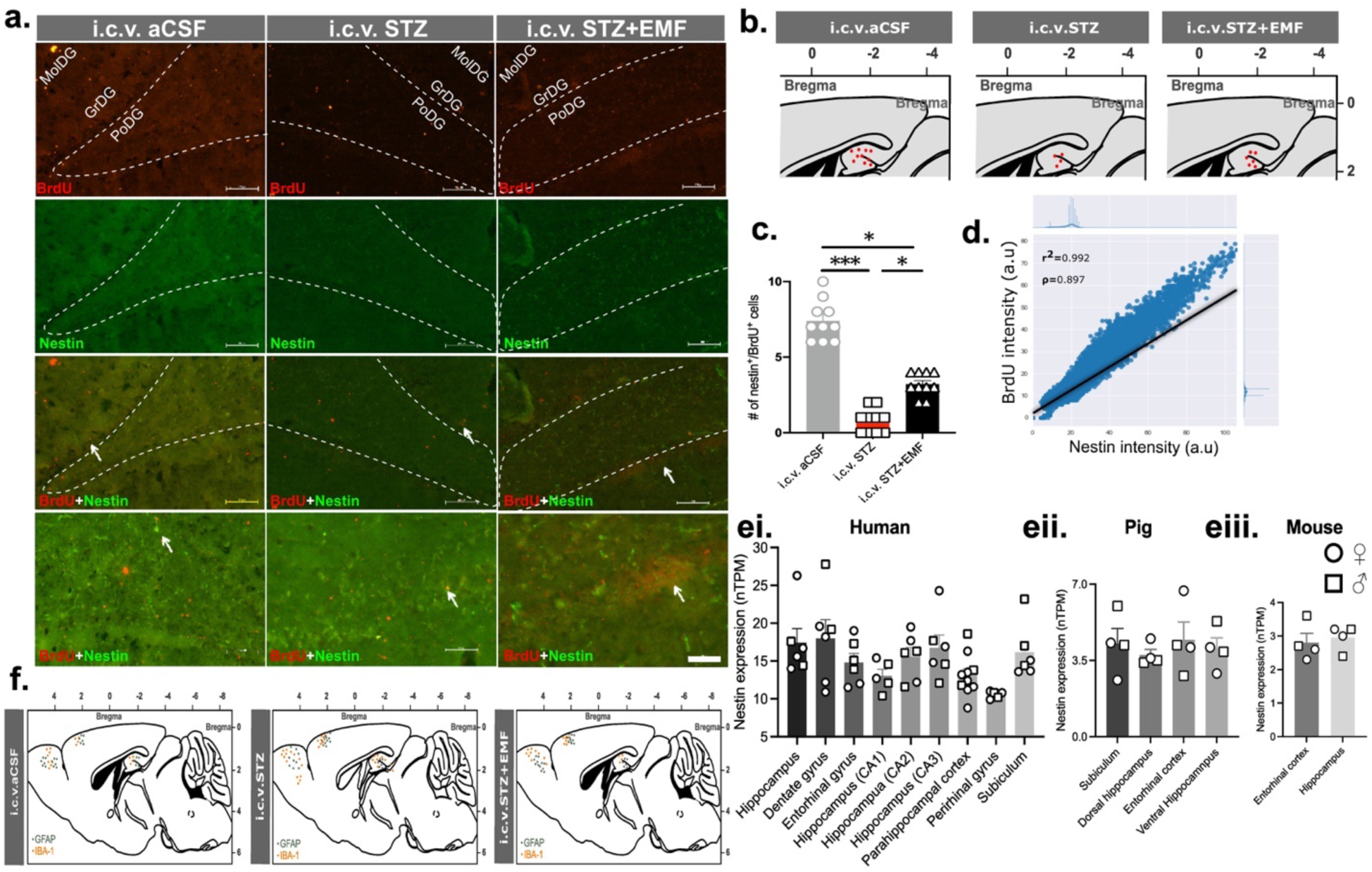
Genesis of new-born neurons in the DG layer of hippocampus upon acute exposure to ELF-MF in STZ injected rats; representative images of the coronal sections from the three groups probed with anti-BrdU and anti-nestin antibody with low and high magnifications with the arrow marking the dual positive cells colocalised in the layers of DG in hippocampus (a); dual positive expression profile plotted across cartesian plane of atlas (b); counting of dual positive cells in the three layers of dentate gyrus (c); scatterplot of BrdU vs nestin intensity with regression line and Pearson correlation co-efficient as measured by just another plugin for colocalization (JACOP) in the i.c.v.STZ+EMF group (d); nestin RNA expression as transcript per million from the mammalian adult neurogenesis gene ontology (MANGO) database(Overall *et al*., 2012b) in human, pig and mouse in two sexes indicated with different markers (empty spherical= male/ empty square= female, ei-iii); representative gliosis profile of the three groups investigated as examined by probing sections with anti-GFAP and anti-IBA-1 antibody for the brain areas like olfactory bulb, frontal cortex and hippocampus (f); data is represented as histogram with individual data points; n=3 for colocalization experiments with different fields as individual data points; p≤0.5 is considered as significant; one-way ANOVA was used for the statistical comparison between three groups whereas Pearson’s correlation was performed for correlation between BrdU and nestin intensity in the i.c.v.STZ+EMF group and for normality check we’ve used Shapiro-Wilk test; no statistics were applied for the nestin RNA expression profile shown for different brain areas of human, pig and mouse from the MANGO database; scale = 50 micron.

Counting of BrdU and nestin +ve cells in the dentate gyrus layer in three group of animals revealed an interesting increase in dual +ve cells in i.c.v.aCSF group (p=0.006) and i.c.v.STZ+MF group (p=0.031) as compared to STZ treated group after 14 days of ELF-MF stimulations. The expression profile in cartesian plane also showed the same alteration confirming the gain of function of nestin postive granular cells in proliferative phase (**Fig**. 3.d-e).

Next, in i.c.v.STZ+MF group after 14 days of non-invasive brain stimulation we found Parson’s correlation (ρ) coefficient was 0.839, and overlap coefficient was 0.99 for BrdU vs nestin signals. The Mander’s coefficient for influence of BrdU signal on nestin was 1.0 whereas the opposite influence was 0.99. The coefficient of determination in regression was r^2^= 0.994. Together counting and cytofluorimetry results suggest strong influence of extremely low frequency magnetic field on upregulation of dual positive signal for BrdU and nestin in dentate gyrus (**Fig.** 3.f).

## Discussions

Alzheimer’s disease (AD) is the most common form of neurodegenerative disorder diagnosed with cognitive dysfunction, primarily due to loss of synapse (DeKosky & Scheff, 1990; Terry *et al*., 1991; Viana Da Silva *et al*., 2019; Taddei & E. Duff, 2025). Though Braak staging based on tau accumulation showed entorhinal cortex to be affected initially, but these synaptic dysfunction is reported to be evident in hippocampus at early stage tightly associated with the cognitive impairments in human patients (Braak *et al*., 2006; Padurariu *et al*., 2012). Dentate gyrus (DG), acts as gateway of neuronal circuitry from layer II of entorhinal cortex to CA3 of hippocampus, and most importantly the mossy fiber is known to process and consolidate memory, with highly plastic changes in molecular and structural organization (Andersen *et al*., 2006; Rebola *et al*., 2017). In the present study, we have reported loss of spatial memory, accompanied by reference memory as measured with MWM and EPM beahviour in i.c.v. STZ treated animals at 3mg/kg of body weight after an incubation period of 14 days. Apart from the increase in the escape latency, time spent in other quadrants except goal quadrant was reflecting these changes, however this was further evident with the less line of crossing in the goal quadrant/ platform zone with thigmotaxic behavior in the streptozotocin treated animals (supplementary **Fig.** 2b). On contrary, after exposure to extremely low frequency magnetic field (ELF-MF) for 14 days we found improvement in the latency and time spent in goal quadrant in MWM test, further transfer latency to the enclosed arm was also found to be reduced in i.c.v.STZ+MF group. Lieu et al., 2015 showed that exposure to 50Hz, 400μT, 24 hour a day for 60 days causes partial improvement in the spatial memory with reduction in Aß deposition in hippocampus of animals simultaneously injected with Aß_1-42_ (5ul/site, single) and D-galactose (i.p., 50mg/kg) (Liu *et al*., 2015). In another study hippocampal injury induced by trimethyltin chloride was protected by ELF-MF stimulation in a customized coil with improvement in cognition, which is associated with adult neurogenesis as shown by BrdU and NeuN (Sakhaie *et al*., 2017). However, the exact intensity or frequency of the magnetic field stimulation was not mentioned in the study. Further, in an study immediately before measurement of spatial memory using MWM test magnetic field exposure to a 50 Hz 8 mT, but not 2 mT magnetic fields for 20 min immediately after training impaired retention performance (Jadidi *et al*., 2007). This study is crucial to indicate that apart from frequency increment in intensity of magnetic field can cause damage to the neuronal circuitry. However, a study with quite similar magnetic field parameter as used in present investigation, 50 Hz, 100µT produced by a circular coils, revealed exposure for chronic time period (90days) for 2hr/day causes no change in the paired pulse ratio in perforant pathway and DG layer of hippocampus. Interestingly, high frequency stimulation causes rise in EPSP of perforant pathway as well as DG granular cells that has been maintained in ELF-MF stimulation group, suggesting induction of long term potentiation in the circuitry mediated by synaptic plasticity (Komaki *et al*., 2014).

Neurodegeneration, synaptic dysfunction in AD is accompanied by neuroinflammation caused by astrocyte as well as microglia (Morales *et al*., 2014; Ransohoff, 2016; Rodriguez-Vieitez *et al*., 2024). I.c.v. STZ induced animal model is also reported to have reactive astrogliosis/ microgliosis with a pattern dependent on dose, incubation period, laterality of injection etc. (Rai *et al*., 2013; Nazem *et al*., 2015; Rostami *et al*., 2017; Ghosh *et al*., 2020; Liu *et al*., 2022). The reason behind this glial aggravation is suggested to be associated with insulin resistance in the central nervous system (Nazem *et al*., 2015).

In the present study, it was evident that there was increased gliosis in both astrocytes and microglia in brain areas like olfactory bulb, frontal cortex and hippocampus after two weeks of i.c.v. STZ injections. Our histometric examination in dentate gyrus of hippocampus also showed a neurodegenerative change as depicted by thinning of granular layer. Interestingly, extremely low frequency magnetic field stimulation of 17.96µT for 2h/day for 14 days improved neuroinflammation as well as the neuronal loss seen in DG. The expression profile of GFAP and IBA-1 signal showed a reduction in volume of inflamed tissue. In a recent work, chronic exposure to 50 Hz ELF-MF at 1 mT for 8 h/day, 5 days/week for 12 consecutive weeks showed no alteration to the GFAP, S100β, phosphorylated tau as well as Aβ expression in hippocampus of aged mice, which suggests ELF-MF does not aggravate aging and associated neuroinflammation, or promote pathological pathways involved in the initiation of AD (Hadzibegovic *et al*., 2025). In a compression model of spinal cord injury, authors studied 20µT intensity with altering frequency at a specific pattern causing pulsed ELF-MF revealing improvements in astrogliosis and microgliosis nearby the site of injury (Goldshmit *et al*., 2022). In another study on 2015, ELF-MF stimulation at 50 Hz, 1 mT in animal model of amyotrophic lateral sclerosis and AD showed protection of reactive gliosis (Liebl *et al*., 2015). Improvements in the glial activity in these studies are also associated with improvements in cognitive function. However, the pattern of stimulation i.e. pulses, and intensity varied across the studies.

Neurodegeneration and neuroinflammation in AD brain are often related to its oxidative balance in the milieu. Previous studies reported elevation of pro-oxidative agents over anti-oxidative agents, only after one week of i.c.v. STZ treatment at doses like 2.5mg-3mg/kg of body weight (Sharma & Gupta, 2001; Mehla *et al*., 2013; El Sayed & Ghoneum, 2020). We’ve seen an increase in SOD1 enzyme activity with reduction in GSH in frontal cortex and hippocampal tissue homogenates in i.c.v. STZ group. This could explain the neurodegenerative changes and glial aggravation observed.

In i.c.v.STZ+MF group after magnetic field stimulation for 14 days we found SOD1 and GSH was restored back in frontal cortex however the hippocampal GSH was not replenished. This suggests ELF-MF treatment in our study could partially improve the redox balance. In a study, authors exposed animals to 7 mT, 40 Hz, either 30 or 60 min/day, 10 days. Results showed partial improvement only in the 60min experimental design as compared to that of 30min counterpart (Ciejka *et al*., 2011). In this study authors mentioned about the adaptation to the external stimulation that may be the reason for beneficial effects in 60min exposure protocol. However, this is important to note that the intensity being chosen for the study in quite high than that of present study. Further, 128 mT, 1 h/day exposure for 30 consecutive days to ELF-MF radiation is shown to improve SOD, GPx activity as well as catalase and lipid peroxidation in frontal cortex as well as partially in hippocampus (Amara *et al*., 2009). Though in this study authors used an static magnetic field. In an animal model of global ischemia, 50Hz and 0.5mT intensity for seven days shoed an reduction in SOD activity, nitric oxide, superoxide production and lipid peroxidation (Balind *et al*., 2014). The results from the prvious studies on the effect of ELF-MF on the oxidative stressors are contradictory. The primary reason, could be the intensity and the duration of the ELF-MF stimulation which could directly influence the redox balance. Further, the procedure for animal sacrifice and tissue collection, more specifically post mortem tissue collection delay could influence the result.

In 2018 Bassani and colleagues examined adult neurogenesis in i.c.v. STZ treated animals. Where they have used neuroinflammation marker GFAP and IBA1, and neurogenesis endogenous markers DCX, KI67, NeuN even with thymidine analogue BrdU in multiple experimental designs. Their observation revealed an increased expression of GFAP, IBA-1 in cortical as well as sub-cortical areas at 3mg/kg dose of STZ after 7 days of injection. On the other hand, DCX, KI67 and NeuN reactivity was reduced in STZ treated animals. BrdU expression Along with KI67 was reduced after 3 weeks of i.c.v. STZ injection (Bassani *et al*., 2018a). It is important to note that in this study authors have used NaCl (0.9%, w/v) as vehicle of STZ, and most importantly the biomarkers used for neurogenesis mainly tag the neurons either at intermediate or mature stage of differentiation. I the present study, we have specifically, wanted to tag the cells that has immature i.e. the nascent stage of the NPSCs cells and that’s why after a rigorous cross-check of the available markes in MANGO database we chose to see the expression of nestin. Nestin is an intermideate filament protein that expresses in multiple cell types from muscle to neuron and depending upon the stage of the cell function could be varied. In the development stage this gene is mainly expressed and can modulate the fate of neuroepithelial cell and its stemness (Herrmann & Aebi, 2000). Interestingly, during embryogenesis, nestin is expressed in migrating and proliferating cells, whereas in adult tissues, nestin is mainly restricted to areas of regeneration (Wiese *et al*., 2004). In a review in 2018 authors has explained potential of nestin as a candidate molecule for the stemness in neural tissues in preclinical models (Bernal & Arranz, 2018). In this study, primary counting of BrdU^+^/Nestin^+^ cell in the granular layer of hippocampus showed an increase in the magnetic field stimulated group, however, the results are also reflecting that the dual positive cell counts were not back to the sham stimulation proxy group i.e. i.c.v.aSCF+EMF. Further, correlation between two signals in magnetic field stimulation group also revealed an tight correlation with similar range of influence of two signals with respect to other as measured with Mander’s co-efficient. Most importantly the behavioural results are supporting our colocalization analysis.

There are many studies available where authors has tried to investigate the effect of magnetic field on the learning and memory with respect to its influence on neurogenesis using multiple tools and experimental approaches (Cuccurazzu *et al*., 2010; Podda *et al*., 2014; Sakhaie *et al*., 2017; Gao *et al*., 2021b; Son *et al*., 2023). Though the intensity and the duration of the magnetic field stimulation were different but the frequencies used in the aforementioned studies were 50Hz and in all the studies, authors noted significant degree of adult neurogenesis in the magnetic field stimulated group. In all of these studies the neurogenesis biomarkers been studied are NeuN or DCX along with or separately proliferative marker BrdU to get holistic picture of the proliferative newborn neurons in the hippocampus DG subfield after ELF-MF stimulation. NeuN and DCX are genes that expressed either at mature (late) or intermediate (middle) stage of neuronal differentiation from NPSCs pools. Hence, the major difference available studies and present one is that in present study we specifically wanted to get idea about NPSCs at early (immature) stage in hippocampus, i.e. before its incorporation to the neuronal circuitry.

In a more recent work authors has used tri-axial square Helmholtz coil and examined effect of magnetic field of 39.4 ± 3.6 μT, which was compared with geomagnetic field range stimulation i.e. 0.17 μT. Results showed impairment in the adult neurogenesis in the dentate gyrus after 39.4 ± 3.6 μT magnetic field stimulation, interestingly it was rescued by 0.17 μT stimulation (Zhang *et al*., 2021). It is noteworthy that in this study authors exposed animals for 5-12 weeks in the simulation parameters mentioned. This suggests that the effects seen in the non-invasive brain stimulation is dependent upon the stimulation parameters.

Taking together all the findings in the present study, we can conclude that acute extremely low frequency (50Hz) magnetic field stimulation at low intensity (17.96µT) with 2hr/day for 14 days causes improvement in spatial and reference memory in i.c.v.STZ treated animals, which could be mediated through improvement in redox balance and neuroinflammation with special emphasis on improvement in adult neurogenesis.

## Author Contributions

AK conducted the experiments collected the raw data for behavioural, biochemical and histological staining; AR conceived the idea, assisted AK with experiment planning, conducted post-hoc imaging analysis and wrote the manuscript; VK conducted immunostaining and performed of immunostaining imaging; SJ supervised for experimental protocols and guided for experiments; JK and YKG supervised with MWM experiments.

## Statement and declaration

### Conflict of interest and ethical statement

Authors in this article claims no potential conflict of interests. Experiments were conducted following guidelines of Institutional Animal Ethical Committee at All India Institute of Medical Sciences, New Delhi vide 12/IAEC/2017.

### Data availability

Raw data analyzed in the present study will be available with request to lead contact.

## Acknowledgements

We acknowledge Mr Sanjeev Beniwal, Department Physiology for assistance in behavioral experiment and Dr Priyanka Kumari, Department of Physiology, All India InsÇtute of Medical Sciences, New Delhi for assistance in histology.

