## Supplementary file for "Non-invasive brain stimulation protects cognitive impairment in i.c.v.STZ injected rats: role of adult neurogenesis"

**Title:**

**Affiliations:**

**Magnetic field stimulation setup:**

In order to give whole-body exposure of magnetic field of ≈17.96μT at a frequency of 50Hz we have customized Lee-Whitting coil in our laboratory at All India Institute of Medical Sciences, New Delhi, India. The structural design of Lee-Whitting coil consists of four circular coils with same diameter with an ampere turn ratio of 9:4:4:9, placed at spacing of -.4704, -.1216, +.1216, +.4704 with respect to the center of the coil (Kirschvink, 1992). This gives the chamber the capability to generate a uniform magnetic field in the center zone ≈6cm (Supplementary **Fig. 1a**). The uniformity of the magnetic field in this circular coil can be calculated by the following formula

We have customized specific plexiglass partial restrainer, with a minute range of flexibility to move extremities of the animals being exposed to the magnetic field during stimulation regime (**Fig. S1b**). The room chosen for this exposure protocol is cross checked for external magnetic field, and this procedure is repeated several times during the whole experiment. This was done to ensure no/ less influence of external magnetic field. Further, we have recorded the magnetic field generated by the system in three coordinates (X, Y, Z-plane/ axis). Our recordings further validated our claim of having a magnetic field intensity of 17.96μT that has been used in the present study (supplementary **Fig. 1d.**). During this recording we have placed the sensor at least three different y-plane at an interval of 2cm to ensure the calculation of the recording cover all the area, where animals were placed during the actual study.

To not have influence of the partial restrainer, on the behavior of the animals we have habituated animals in the partial restrainer for at least 4-5 days before actual exposure. Next, the duration of the exposure to the magnetic field has been chosen from previous studies on not only STZ-induced AD animals from the laboratory but also from the previous studies in other animal models like 6-OHdA induced animal model of Parkinson’s disease and studies on spinal cord injury both transection model as well as with New York University impactor device (Bose *et al.*, 2023; Kaur *et al.*, 2024).

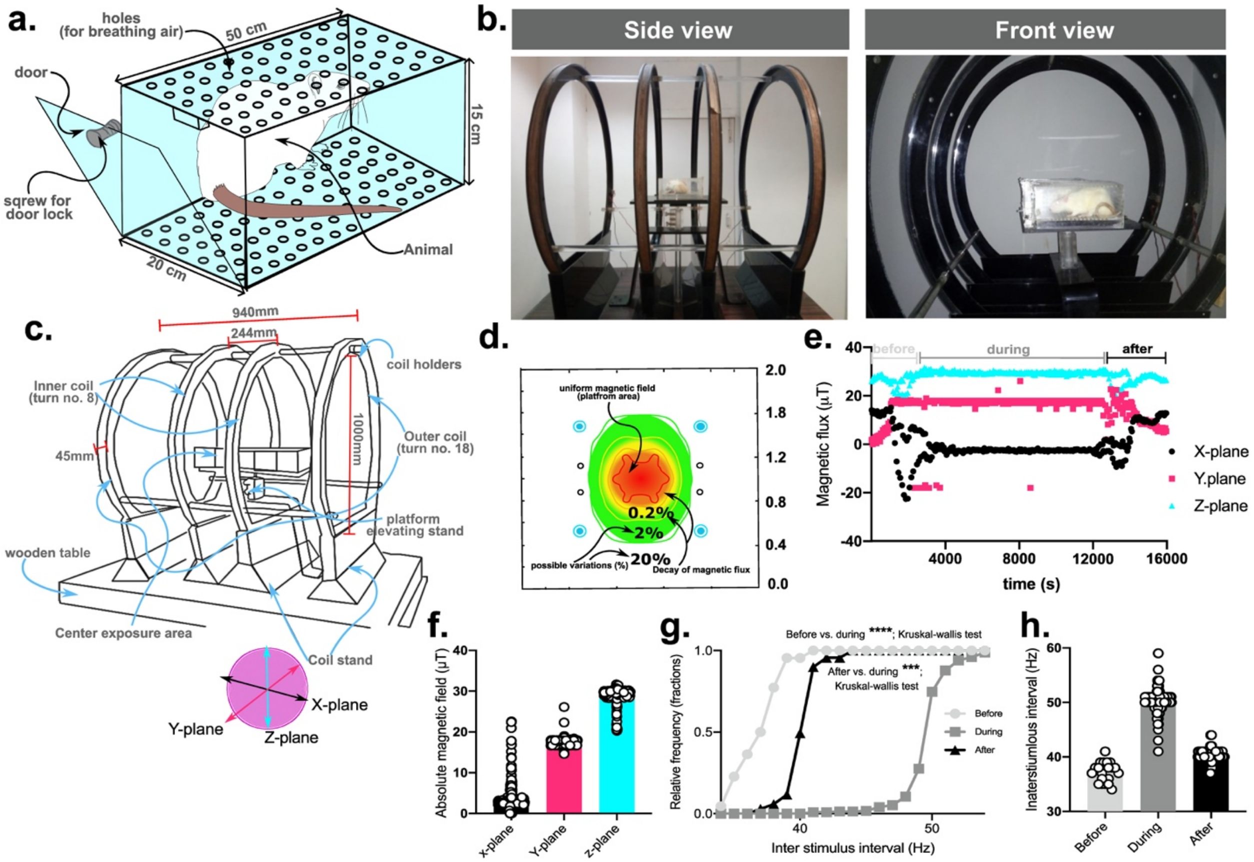


**Supplementary figure 1.: Extremely low frequency electromagnetic field exposure setup;** illustration of partial restrainer used for animals to be kept in the magnetic field chamber during exposure (a); side and front view of the actual Lee-Whitting coil/ modified Helmholtz coil used to expose the animals in the magnetic field regime in the study (b); blue print of the coil setup with exact dimensions of its coil and other associated accessories (c); magnetic flux distribution at 2X2 meter area modified from (Kirschvink, 1992) with colour coding for ‘hot’ to ‘cold’ spot representing the uniform distribution (d); real-time recording of magnetic field for 4.4hr with before and after switching on the electromagnetic coil marked as ‘before’, ‘during’, and ‘after’ with three axes (e); absolute magnetic field at different planes (f); relative frequency distribution and average inter stimulus interval i.e. frequency before-during-after magnetic field stimulation (g); data is represented as either scatter plot for 4hr magnetic field recording and histogram with individual data points for other variables; Kolmogorov-Smirnov test was applied for relative frequency distribution for inter stimulus interval; no statistics were performed for other parameters.

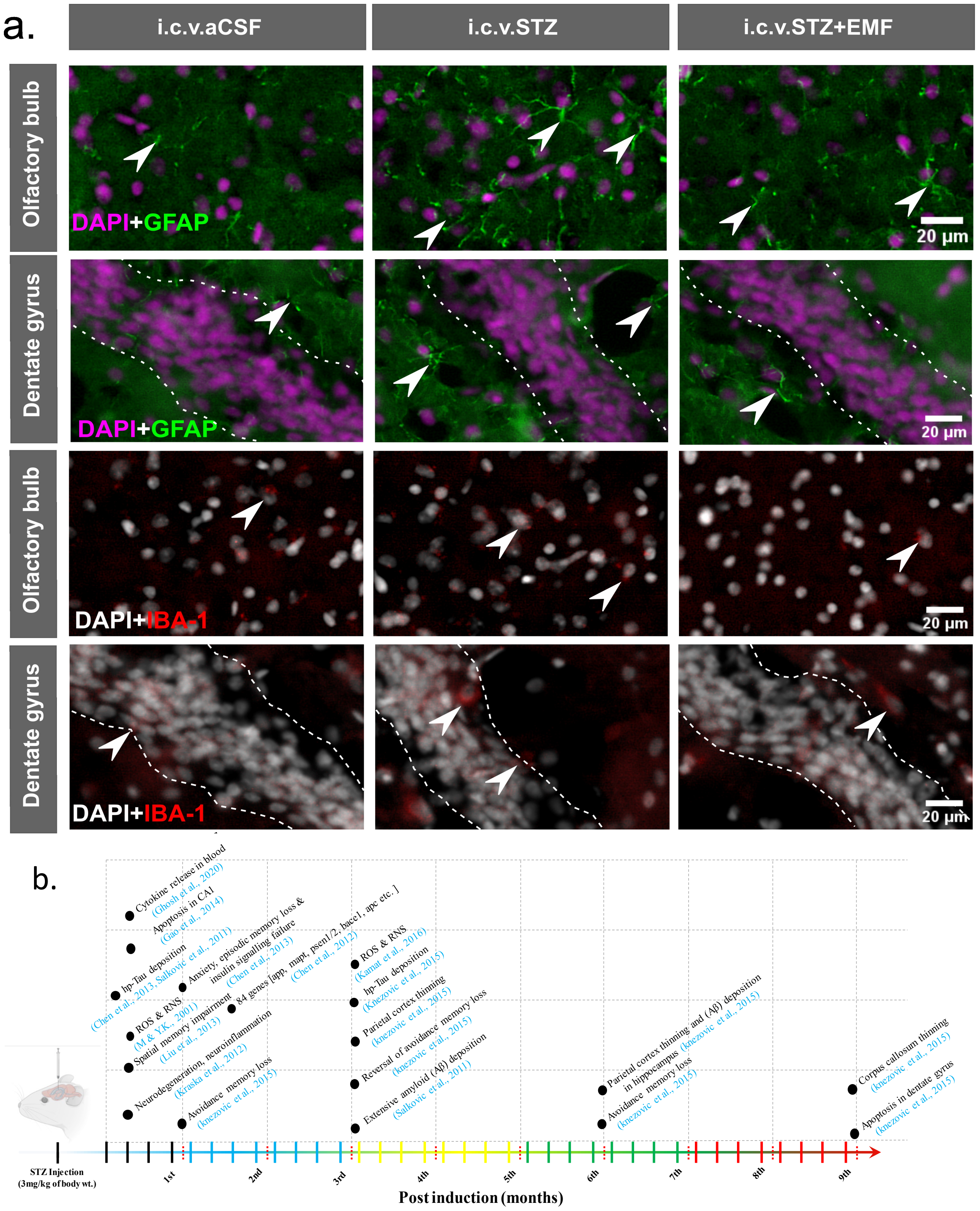


**Supplementary figure 2.: Amelioration of gliosis upon exposure to extremely low frequency magnetic field;** representative images of GFAP (in green) and IBA-1 (in red) sub cellular expression in olfactory bulb and dentate gyrus with arrow heads denoting the positive cells; all the sections were counterstained with DAPI, however they were pseudo-colored ‘purple’ in GFAP tiles and ‘grey’ in IBA-1 tiles (a); chronological order of effect reported in the previous reports(Sharma & Gupta, 2001; Salkovic-Petrisic *et al.*, 2011; Chen *et al.*, 2012, 2013; Kraska *et al.*, 2012; Liu *et al.*, 2013; Gao *et al.*, 2014; Knezovic *et al.*, 2015; Kamat *et al.*, 2016; Ghosh *et al.*, 2020) in the i.c.v.STZ treated animal model for Alzheimer’s disease note majority of the effects were being reported within two weeks post-injection of STZ (b); scale = 20 micron.
